# Evolutionary phenome-genome analysis of cranial suture closure in mammals

**DOI:** 10.1101/2020.06.15.148130

**Authors:** Borja Esteve-Altava, Fabio Barteri, Xavier Farré, Gerard Muntané, Juan Francisco Pastor, Arcadi Navarro

## Abstract

Cranial sutures are growth and stress diffusion sites that connect the bones protecting the brain. The closure of cranial suture is a key feature of mammalian late development and evolution, which can also lead to head malformations when it occurs prematurely (craniosynostosis). To unveil the phenotypic and genetic causes of suture closure in evolution, we examined 48 mammalian species searching for (i) causal links between suture patency, brain size, and diet using phylogenetic path analysis; and (ii) instances of genome-phenome convergence amino acid substitutions. Here we show that brain size and the anteroposterior order of ossification of the skull are the two main causes of sutures patency in evolution. We also identified three novel candidate genes for suture closure in evolution (HRNR, *KIAA1549*, and *TTN*), which have never been reported in clinical studies of craniosynostosis. Our results suggest that different genetic pathways underlie cranial suture closure in evolution and disease.

## INTRODUCTION

Cranial sutures separate the bones of the skull and function as sites of bone growth and stress diffusion (Herring, 2008; Opperman, 2000). They are necessary to develop a healthy, functioning head in mammals. Interestingly, while many sutures remain open through life, some cranial sutures will naturally close by turning into bone. The closure of suture is a key feature of the mammalian skull late development (**Figure 1**), growth, functioning, and evolution (Cray et al., 2014; Oh et al., 2017; Roston & Roth, 2019). However, a premature closure of sutures (craniosynostosis) can also lead to head malformations in newborns (Cohen & MacLean, 2000).

**Figure 1.**
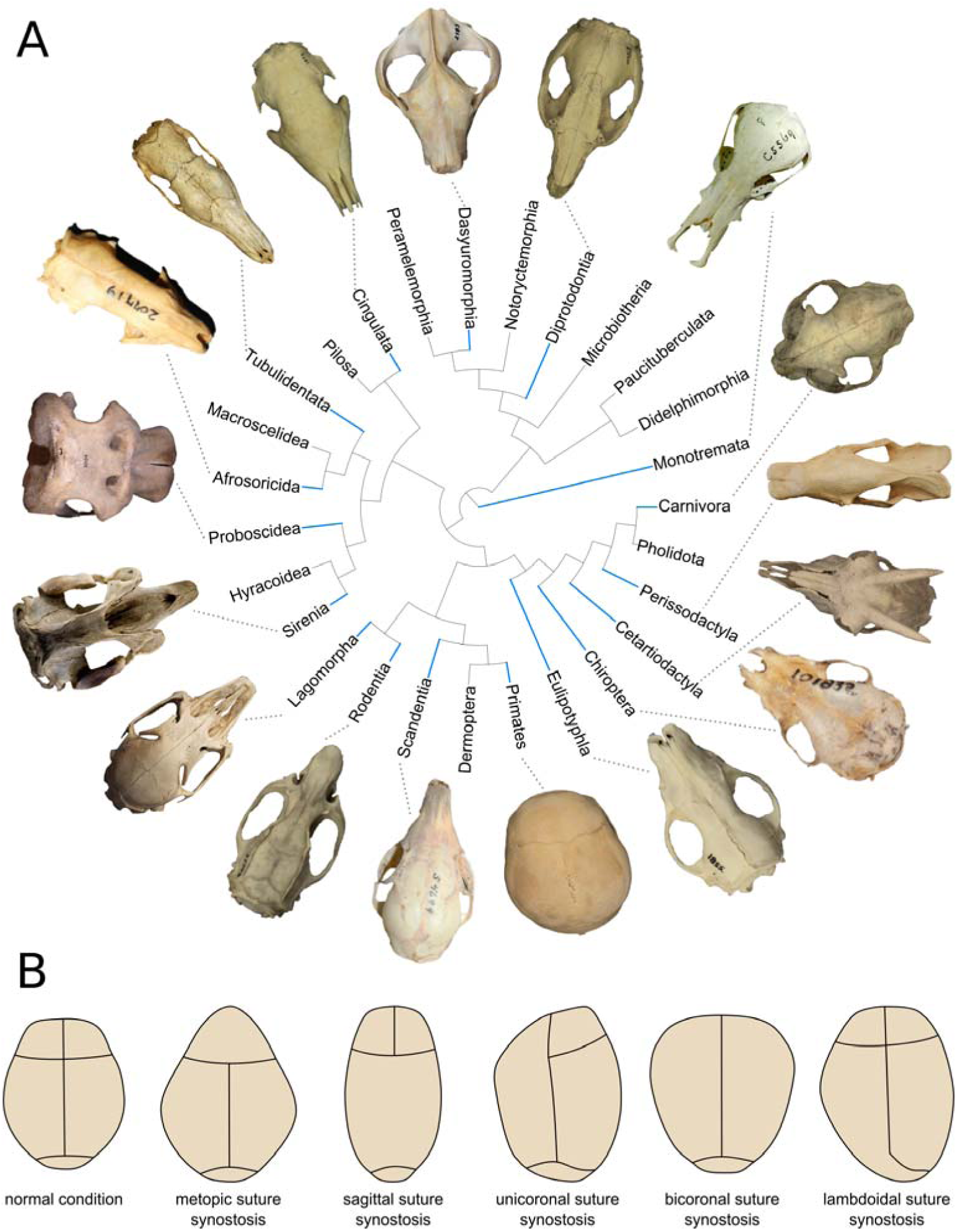
Cranial suture close in evolution and disease. (**A**) Mammalian orders studied (blue lines), with representative skulls in dorsal view showing the differences in patency for the metopic, coronal, and sagittal sutures. Some mammals close their cranial sutures after birth, and we see them closed in adults, while others keep their sutures open through life or until an old age. (**B**) Outlines showing the consequences in the shape of the skull by a premature suture closure in humans, a condition called craniosynostosis.

In mammals, cranial sutures closure evolves in coordination with the rest of the body. In fact, suture closure is positively correlated with skull size and body size (Bärmann & Sánchez-Villagra, 2012; Wilson & Sánchez & Villagra, 2009). Cranial sutures and the brain are also tightly integrated by physical contiguity and shared signaling pathways during development (Lieberman, 2010; Richtsmeier & Flaherty, 2013). Thus, changes of brain size could affect suture closure in evolution. Likewise, diet can cause suture closure as a secondary effect of the mechanical loads generated during feeding (Rafferty & Herring, 1999; Sun et al., 2004; Rafferty et al., 2019; Byron et al., 2004). Biomechanical studies show that compression on a suture creates an environment that favors suture closure (e.g., enhances osteogenesis, narrows suture space, immobilizes bones), whereas tension favors suture patency (Herring, 2008). Therefore, we could expect suture closure to be a by-product of the evolution of these traits, particularly brain size and diet. To our knowledge, there are no studies on the impact of these phenotypic traits to suture closure in evolution.

Anatomical constraints can also bias which sutures close and which remain open (Moss, 1975; Lieberman, 2011; Esteve-Altava & Rasskin-Gutman, 2014; Rasskin-Gutman & Esteve-Altava, 2018). Theoretical models predict that the whole arrangement of sutures in the skull—as a network—acts itself as an anatomical constraint that influences which sutures will close and which ones will remain open (Esteve-Altava et al., 2017), for example, by directing the signaling pathways that promote osteogenesis through mechanosensors (Khonsari et al., 2013; Katsianou et al., 2016). Additionally, the timing of ossification of skull bones, from face to vault (Koyabu et al., 2014), and of suture closure, from vault to face (Rager et al., 2014), have also the potential to explain biases in suture closure, since one suture closure may influence the closure of a neighboring suture.

The genetic causes of suture closure in evolution remain largely unknown. Most of our knowledge comes from medical studies of genetic syndromes causing premature suture closure in humans (Lattanzi et al., 2017; Morriss-Kay & Wilkie, 2005; Poot, 2019; Twigg & Wilkie, 2015; Wilkie et al., 2017) and animal models (Cornille et al., 2019; Grova et al., 2012). These studies have revealed a complex network of genes involving many signaling pathways (e.g., *BMP*/*TGF*-β, *FGF*, and *WNT*). However, about 80% of craniosynostosis cases are nonsyndromic: they typically affect only one suture and are not associated with other body malformations (Dempsey et al., 2019; Garza & Khosla, 2012). There is little information on the genetic causes of nonsyndromic craniosynostosis (Sewda et al., 2019). Evolutionary genomics offers a powerful tool to explore the genetic causes of natural variation (Alföldi & Lindblad-Toh, 2013; de Magalhães & Wang, 2019; Smith et al., 2020), as shown by studies on skull shape evolution (Roosenboom et al., 2018) and marine adaptations (Foote et al., 2015; Zhou et al., 2015). Evolutionary studies have shown that some of the genes regulating suture closure (e.g., *BMP3, MSX2, RUNX2*) have evolved under positive selection in humans and other primates (Green et al., 2010; Magherini et al., 2015; Twigg et al., 2015; Wu et al., 2010, 2012), which suggests that the same processes favoring suture closure at evolutionary scale could be causing craniosynostosis conditions. Similarities of potential genetic factors and phenotypes between skull evolution and craniosynostosis (e.g., fewer bones, same sutures frequently affected, related shape changes) could indicate that analogous mechanisms underlie suture closure in evolution and disease (Esteve-Altava et al., 2017; Richtsmeier, 2018; Richtsmeier et al., 2006).

Here we assessed the evolutionary factors determining the closure of the metopic, coronal, and sagittal suture in mammals. To this end, we analyzed the cranial anatomy of 48 species of mammals, for which their reference genomes were aligned at UCSC (100-way) and for which there was information on their diet and brain mass in the literature. First, we tested 12 alternative evolutionary hypotheses of how brain size, diet, and developmental constraints may determine suture closure in evolution. Then, we looked for convergent amino acid substitutions in multiple-sequence alignments of proteins, comparing species with sutures closed or open. Our hypotheses are that (1) brain size, diet hardness, and constraints, together, determine suture patency and closure in evolution; (2) species with a given suture closed will share mutations in the same key genes that will be absent in closely related species with the same suture open; (3) these genes will be enriched in biological functions relevant to cranial suture formation and maintenance, brain growth, and biomechanical performance; and (4) they will overlap with genes previously associated to craniosynostosis in clinical studies.

## RESULTS

The patency of the metopic, coronal, and sagittal sutures varied in the sample set of mammals analyzed, with some taxonomic groups showing a consistent pattern of closure for some sutures (**Figure 2**). We used the frequency of suture patency (i.e., specimens with the suture open/total specimens) to infer the causal links between suture closure and other phenotypic traits of interest in a phylogenetic path analysis. The high degree of conservation of suture patency within species enabled us to categorize sutures phenotype, as either open or closed, and to carry out a search for convergent amino acid substitutions in the protein-coding genes.

**Figure 2.**
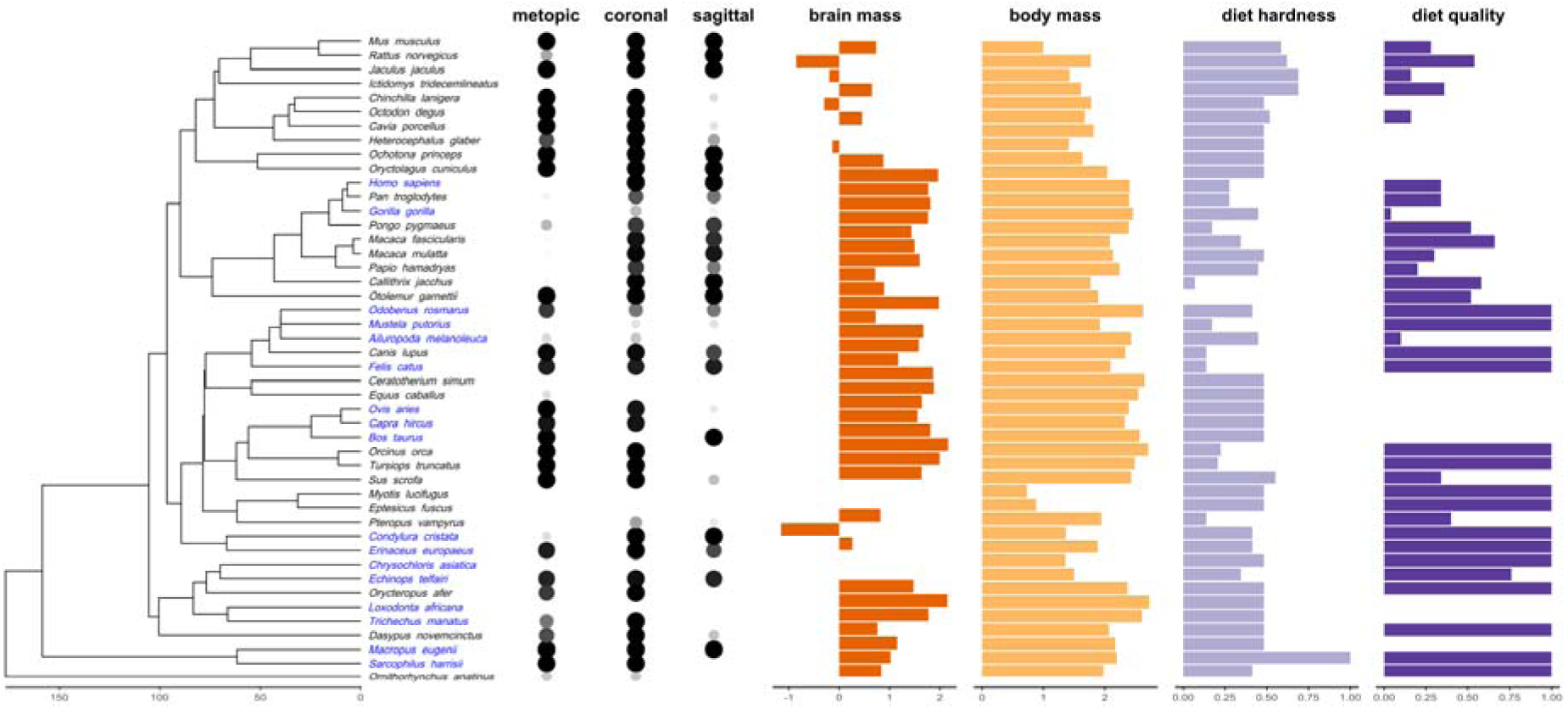
Variation in suture patency and potentially related traits in mammals. In blue, species used for the pair-wise convergent amino acid substation analysis. Note that different pairs of species were used for each suture and that only suture patency below/above the threshold of 0.25/0.75 were considered. Dot size and gray scale of suture patency ranges between 0 to 1, corresponding to 0% to 100% of specimens with the suture open in our sample. Brain and body mass shown as log-transformation of weight in grams. Diet hardness and quality are normalized. See **Supplementary File 1** for the exact values of each variable.

### Suture patency and life-history traits evolution

The best supported model for the patency of the metopic, coronal, and sagittal sutures in evolution agrees with the hypothesis of an anteroposterior direction in sutures’ closure following the timing of skull ossification and with the influence of brain size after correcting for body size (CICc = 56.9, *p-value* = 0.196, *w* = 0.829). (N/A: CICc is a modified version of AICc for path analysis, significant p-values, < 0.05, mean that the model is rejected, see *Methods*). **Figure 3** summarizes the results of the phylogenetic path analysis. The best model includes mid-to-high effects of one suture on another in an anteroposterior direction and low-to-mid effects of brain size on sutures. Larger brains (after correcting for body size) tend to favor the maintenance of the coronal and sagittal sutures open, and to a lesser extent, the closure of the metopic. In contrast, diet quality has a negligible positive effect on brain size. Regardless of the support of each model (ΔCICc), only those hypotheses that included a causal relation between sutures and brain size were supported by evidence (*p-value* > 0.05). See **Supplementary File 1** for details.

**Figure 3.**
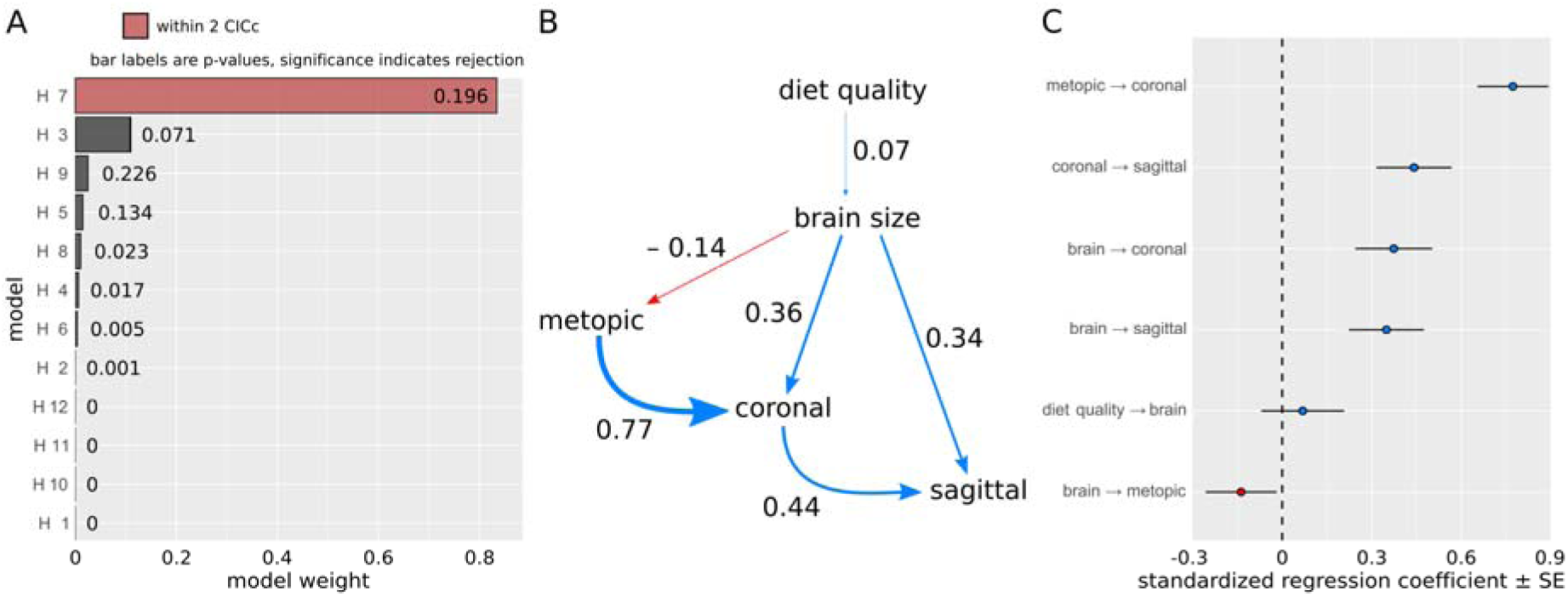
Summary of the phylogenetic path analysis. (**A**) Comparison of statistics for each model. There is only one best supported causal model (i.e., within 2 CICc and *p-value* > 0.05). (**B**) The best model shows a causal relation for cranial sutures patency following an anteroposterior direction, with larger brain sizes causing the sutures coronal and sagittal to remain open and the metopic suture to close at an evolutionary scale. An arrow indicates the direction of the causal relation, its width corresponds to the standardized regression coefficient (i.e., how much the causal variable affects the effect variable), and its color represents a positive (blue) or a negative (red) effect. (**C**) Amount of change (with standard error) produced by causal variables on effect variables for the best supported model.

### Convergent amino acid substitutions in species with cranial sutures closed

We compared the amino acid sequences of 10,922 genes in pairs of closely related species that vary in their suture patency (open *versus* closed). We recovered an aggregate of 6,158 putatively convergent amino acid substitutions (CAAS) in 3,250 unique genes. These genes were mutated in at least one suture, meaning that they were identified in three pair-wise comparisons of closely related species with different suture patency. The number of unique genes found is significantly lower than expected at random (1,000-bootstrap: α^0.05^ = 5,759.8; see *Methods* for more details). We recovered a total of 28 candidate genes that overlap in the three sutures (i.e., identified in nine pair-wise comparisons), which were used to perform set enrichment analyses. This is a significant overlap between the genes identified for each suture compared to that expected if the three sets were independent (*Fold Enrichment* = 15, p-value = 5.04E-25). Out of the 28 candidates, three genes were also internally validated in the whole sample of mammals: Hornenin (*HRNR*), *KIAA1549*, and Titin (*TTN*). We focused our discussion on these three gene and their potential role in suture closure. See **Supplementary File 1** for the complete list of CAAS.

### Functional and pathological enrichment

Only one GO functional enrichment was found for the list of 28 candidate genes shared between the three sutures: a molecular function related to limpid transport (GO:0005319, ER = 20.97, FDR = 0.013). For comparison, we also performed functional enrichment analyses for genes identified in at least two sutures (i.e., found in six pair-wise comparisons). This shows that other well supported GO enrichments for these genes include, for example, biological adhesion (GO:0022610, ER = 2.39, FDR = 0.005) and components of the extracellular matrix (GO:0031012, ER = 2.47, FDR = 0.006). See more details in **Supplementary File 1**. Overall, these results provide little support to our initial working hypothesis that candidate genes would be enriched in functions commonly associated with suture biology.

Moreover, neither of the 28 candidate genes overlaps with craniosynostosis genes as ascertained from the study of human disease. None of the candidate genes is present in the lists of genes from the literature (set 1), from upregulated in nonsyndromic craniosynostosis osteoblast cultures (set 2), or from GWAS of nonsyndromic metopic and sagittal cases (set 3). As a reference, we also estimated the overlap between these three gene sets. One-sided Fisher’s exact tests show that only one overlap for the list of literature genes and the list of GWAS (set 1 vs set 2: Odds Ratio = 1.616, FDR = 0.106; set 1 vs set 3: Odds Ratio = 93.29, FDR = 2.05E-30; set 2 vs set 3: Odds Ratio = 1.08, FDR = 0.807). In contrast to candidate genes, genes associated to human craniosynostosis are enriched in biological functions more related to suture biology (see **Supplementary File 1**). This result rejects our starting hypothesis that candidate genes, with mutations linked to suture closure in evolution, would overlap with genes associated to craniosynostosis.

## DISCUSSION

Our results suggest that cranial suture closure in evolution is regulated by two developmental processes: the order of ossification of skull bones and the growth of the brain relative to body size. In contrast, we found no support for diet hardness (as a proxy for chewing biomechanics) as a cause for differences in suture patency among species. From a genetic point of view, we identified 28 candidate genes for suture closure in evolution, out of which three show the strongest support: *HRNR, KIAA1549*, and *TTN*. These genes have never been causally linked to craniofacial development, suture biology, or craniosynostosis. However, they are expressed in tissues adjacent to cranial sutures, such as the brain and skeletal muscles, which affect suture biology.

### Brain size and ossification timing determine suture patency in evolution

Brain growth is tightly integrated with cranial suture maintenance and closure in normal development and in pathological conditions (Richtsmeier, 2018; Richtsmeier et al., 2006; Richtsmeier & Flaherty, 2013). The traditional idea is that suture closure is a passive consequence of the growth of the brain (Moss & Young, 1960; Moss, 1975), with dura mater triggering a signaling cascade that promotes osteoblast activity and regulates cell proliferation and apoptosis (Opperman, 2000; Spector et al., 2002). Our results support a direct influence of brain size on suture patency in mammals, acquired by an increased brain growth relative to body size. However, each suture responds differently to brain size. Larger brains cause the coronal and sagittal sutures to remain open, while they cause the closure of the metopic suture. Why sutures respond in a different way to brain size is unclear, but it could be a consequence of how the growing brain interacts differently with each of the enclosing bones and sutures (Barbeito-Andrés et al., 2020).

Our evolutionary analysis shows that the patency of a suture depends also on the patency of other sutures, following the most common anteroposterior order of ossification of the skull in mammals (Koyabu et al., 2011, 2014). This result agrees with recent theoretical findings proposing that the organization of the skull, as a network of bones connected by sutures, can bias suture closure (Esteve-Altava et al., 2017; Esteve-Altava & Rasskin-Gutman, 2015). For example, by directing mechano-transduction and morphogenetic signals (Khonsari et al., 2013; Katsianou et al., 2016). If that is true, this means that sutures are not only passive subjects of their underlying functional matrices (Moss, 1975), instead they can constrain each other’s closure. However, the exact relation between the timing of ossification and suture closure during development is still unclear and may depend on other species-specific anatomical constraints. In humans, for example, the later closure of the metopic suture compared to our extinct relatives and other primates is thought to be an adaptive response to pelvic constraints on the birth canal (Falk et al., 2012).

### Mechanical stress and suture closure

Biomechanical studies on vertebrate skulls as disparate as lizards and mammals have shown that cranial sutures relieve strain locally in response to mechanical loads, for example, from chewing (Herring & Teng, 2000; Moazen et al., 2009; Rafferty & Herring, 1999). However, using diet hardness as a proxy, we found no support for an effect of mechanical stress on sutures patency or closure in mammals. This is either because there is no evolutionary relation between them or because diet hardness is not a good proxy for the mechanical stress supported by the skull. If diet hardness is not a good proxy, the most accurate alternative would be to carry out biomechanical studies on every species and measure each suture response individually, for example, using Finite Elements Analysis (Bright, 2012). This would be a feasible albeit challenging empirical work.

### New candidate genes for suture closure in mammals

Following a comparative analysis of protein-coding regions, we identified 28 candidate genes that may have a potential role in determining cranial suture closure in mammalian evolution. Out of this list, three genes—*HRNR, KIAA1549*, and *TTN*—are the most likely candidates, because they also show convergent mutations across the whole sample of mammals. These three genes have never been associated with neither normal nor pathological craniofacial development. Thus, we can only speculate about their relation to suture closure based on indirect evidence, such as the tissues where they are expressed, their biological function, and their relations to other proteins.

*HRNR* encodes for a profilaggrin-like protein that functions as an ion binder for calcium and other metals in different tissues (mostly the skin, but also in the brain), organizing the cell envelope and extracellular keratinization. Although *HRNR* was tentatively reported as a risk factor for craniosynostosis in a study of twins (Rymer, 2015), this result has not been validated (*personal communication*). *HRNR* is not known to play a role in cranial development, but other proteins related to keratinization have been reported to participate in developing calvarial bone and sutures (Atsawasuwan et al., 2013). A total of 59 different mutations were found along the entire protein of *HRNR*, none of which targets a functionally active region for calcium binding. However, *HRNR*-coding region is enriched in methylation sites that undergo modifications during human development from newborn to adult (Salpea et al., 2012), the time when cranial sutures close. We identified 58 convergent substitutions for *HRNR* in our evolutionary study (the most of all candidate genes), which could be interacting with methylation sites that determine cranial phenotypes during postnatal development.

*KIAA1549* encodes for a protein component of the cellular membrane that is highly expressed in the brain. Eleven different mutations were found for *KIAA1549*, none within the transmembrane region. *KIAA1549* has never been associated to craniofacial morphology or premature suture closure, but through a fusion with the B-Raf proto-oncogene (*BRAF*) it has been associated to developing pilocytic astrocytoma, a benign brain tumor (Yamashita et al., 2019). Interestingly, *de novo* mutations in *BRAF* has been recently discovered in patients with isolated sagittal synostosis (Armand et al., 2019; Davis et al., 2019). Whether *KIAA1549* can affect suture development through its effect on *BRAF* is not known.

*TTN* encodes for the largest human protein, a common type of filament present in cardiac and skeletal muscles that is essential for muscle contraction. Thirty-two mutations were found for *TTN*, none within its active sites. Although *TTN* has not specifically been associated with craniofacial development or dysmorphologies, we known that head muscles activity is necessary for the correct formation and maintenance of cranial sutures (Byron et al., 2004; Herring & Teng, 2000; Moss & Young, 1960). For example, osteoblast in the sutures respond to muscle tension by increasing the formation of bone (Herring, 2008), which is something also observed in craniosynostosis (Al-Rekabi et al., 2017). We can only speculate on whether small changes in *TTN* proteins modulate head muscles contraction during cranial development, and by doing so, can alter cranial sutures maintenance and closure.

The 28 candidate genes for cranial suture closure in evolution are not enriched in biological functions and cellular components often associated with skeletal development (e.g., osteogenesis, growth factor binding, cell proliferation), which are those dysregulated in pathological cases of suture closure (Rojas-Peña et al., 2014). Instead, we found that candidate genes are functionally enriched in proteins for the transport of lipids across the membrane. Lipids play an important role in skeletal metabolism, for example, by limiting permeability of the bone surface and by regulating biomineralization through the transport of essential fat-soluble vitamins D and K (Tintut & Demer, 2014). In the context of cranial sutures, a relationship between vitamin D deficiencies (congenital or nutritional) and craniosynostosis has been known for a long time (Imerslund, 1951; Jaszczuk et al., 2016; Wang et al., 2015). This result suggests that mechanisms of cranial suture closure in evolution could evolve through changes in the regulation of vitamin D transport, rather than acting directly on osteological regulation pathways. Finally, for genes differentially mutated in two out of three of the sutures, we did find enrichments for biological adhesion and components of the extracellular matrix, which are essential for the maintenance and closure of cranial sutures (Opperman & Rawlins, 2005; Stamper et al., 2011).

### Are evolution and pathological development decoupled for cranial suture closure?

Candidate genes from our evolutionary analysis show a complete lack of overlap with genes linked to pathological suture closure (Adhikari et al., 2016; Justice et al., 2012, 2020; Rojas-Peña et al., 2014). There are many plausible reasons that could explain this mismatch, from methodological limitations to biological causes. On the one hand, the list of genes compared could be incomplete. This is because either (1) our evolutionary analysis fails to identify candidate genes for suture closure or because (2) we only know a limited number of genes which mutation lead to premature suture closure. The first reason would imply that our approach does not work for this phenotype, because it cannot capture mutations affecting the timing of closure (only the mechanism performing the closure), whereas pathological conditions maybe occur due to mutations affecting timing exclusively (e.g., via ectopic gene expression, Poot, 2019). The second reason would mean that our current knowledge of the genetic origins of nonsyndromic craniosynostosis is limited and, therefore, the lists of genes compared fail to capture the complete genetic landscape of this complex disease (Magalhães & Wang, 2019; Lattanzi et al., 2017). On the other hand, there may be biological reasons that explain this mismatch. As mentioned before, changes in brain growth rates or vitamin D regulation in evolution could be one of such underlying causes, acting differentially for each suture (Barbeito-Andrés et al., 2020).

Another possibility is that evolutionary mechanisms are decoupled from developmental mechanisms commonly disrupted in disease, so that analogous phenotypic changes (i.e., closing a suture) can proceed through different paths. Decoupling evolutionary mechanisms of phenotypic variation from those genetic pathways whose disruption is most likely to be detrimental for the individual could be a way to maintain evolvability without compromising fitness, bypassing pleiotropic or epistatic constraints (Payne & Wagner, 2019). The fact that candidate evolutionary genes for suture closure show no enrichment in any disease set supports this hypothesis. However, it is unclear whether macroevolutionary genetic changes should involve the same loci or mutations uncovered by microevolutionary and clinical studies (Smith et al., 2020). For example, the Runt-related transcription factor 2 (*RUNX2*) is a strongly supported candidate to drive facial morphological and suture closure in human evolution (Adhikari et al., 2016; Magherini et al., 2015) and which mutation causes craniosynostosis (Cuellar et al., 2020; Maeno et al., 2011). However, *RUNX2* takes no part in marsupial craniofacial diversity (Newton et al., 2017). This suggests that different mammalian clades can use alternate pathways to control the exact same phenotypic traits.

### Conclusion

Our study dissected the phenotypic and genetic causes of cranial suture patency in evolution, highlighting developmental and evolutionary factors for suture closure in mammals. From a phenotypic point of view, we found two main factors: (1) brain growth, which was a known cause of suture patency in normal and pathological development, and (2) sutures self-regulation, which was previously suggested only by theoretical models. From a comparative genomics approach, we identified candidate genes involved in lipid transport, cell adhesion, and the formation of the extracellular matrix. The best supported candidate genes to play a role in cranial suture closure in evolution are *HRNR, KIAA1549*, and *TTN*. If validated by additional comparative analyses, and experimentally in model organisms for suture closure (e.g., zebrafish, mice, or rabbit), they could provide new ways to study the genetic basis of suture closure in evolution and disease. To our knowledge, this study is the first attempt to search for the genetic causes of cranial suture closure and associated pathologies at a macroevolutionary scale. Our findings highlight the importance of evolutionary approaches to make new discoveries and test hypothesis on development and disease.

## METHODS

### Sampling

We surveyed an initial sample of 53 species of mammals with multiple sequence alignments of their reference genomes available at UCSC (Kent et al., 2002) and with reliable information on their diet, brain mass, and body mass (Burger et al., 2019). We examined adult skull specimens in vivo and in digital images from museums and online collections with an ID catalog number. A total of 48 species had more than two well-preserved specimens available for examination and were included in the present study. Details for specimens ID, suture patency, life traits, phylogeny, and analysis code are available at https://figshare.com/projects/Cranial_Suture_Closure/81209.

### Suture patency

For each specimen, we coded the state of the metopic, coronal, and sagittal sutures as either open or closed, depending on whether they were visible (patent) or not (obliterated). Ambiguous cases (e.g., when a suture is half closed) were rare and we excluded them from the study. Suture patency was quantified as the ratio of the number of specimens with the suture open to the total number of specimens examined, ranging from 1 (all open) to 0 (all closed). This continuous measure provided an amenable variable to perform the phylogenetic path analysis. We omitted from our analyses the sagittal suture in the orca and the dolphin, because cetaceans never form this suture in the first place due to the expansion of the occipital bone (Roston & Roth, 2019).

To later search for convergent amino acid substitution (CAAS), suture patency was binarized by thresholding it between 0.75 and 0.25. A suture with a patency above 0.75 was counted as open and below 0.25 was counted as closed. A binarization of suture patency was necessary because CAAS comparisons require as input two discrete groups of species (Muntané et al., 2018). Because suture patency is a highly conserved trait, most species ranked well above or below these thresholds. Only 15 sutures out of 144 observations were left uncategorized, and we omitted them when selecting pairs of species to compare their protein-coding sequences.

### Life traits and phylogenetic path analysis

We tested 12 alternative causal models for the closure of the metopic, coronal, and sagittal sutures using a phylogenetic confirmatory path analysis (von Hardenberg & Gonzalez & Voyer, 2013). To this end, we downloaded a calibrated phylogeny for all mammals from TimeTree (http://www.timetree.org/) and pruned off the species not sampled. The analysis was carried using the Pagel’s lambda model of evolution, which is estimated internally by the function *phylo_path* (Bijl, 2018) in R (R Core Team, 2019). **Figure 4** shows the models evaluated. As potential relations we modeled the mutual causation between sutures due to their development (Koyabu et al., 2014; Rager et al., 2014), the effect of hard diet on sutures due to the stress involving in chewing (Herring, 2008; Rafferty et al., 2019; Sun et al., 2004), and the effect of brain size on sutures due to the influence of the brain on the growth of the bones of the vault and the maintenance of sutures (Richtsmeier, 2018; Richtsmeier et al., 2006; Richtsmeier & Flaherty, 2013). Finally, we included the effect of diet quality on brain size as an indirect link on sutures (Aiello & Wheeler, 1995; Allen & Kay, 2012). To reduce the number of variables in the models, body size was included as a corrector for brain size, instead of as an independent variable.

**Figure 4.**
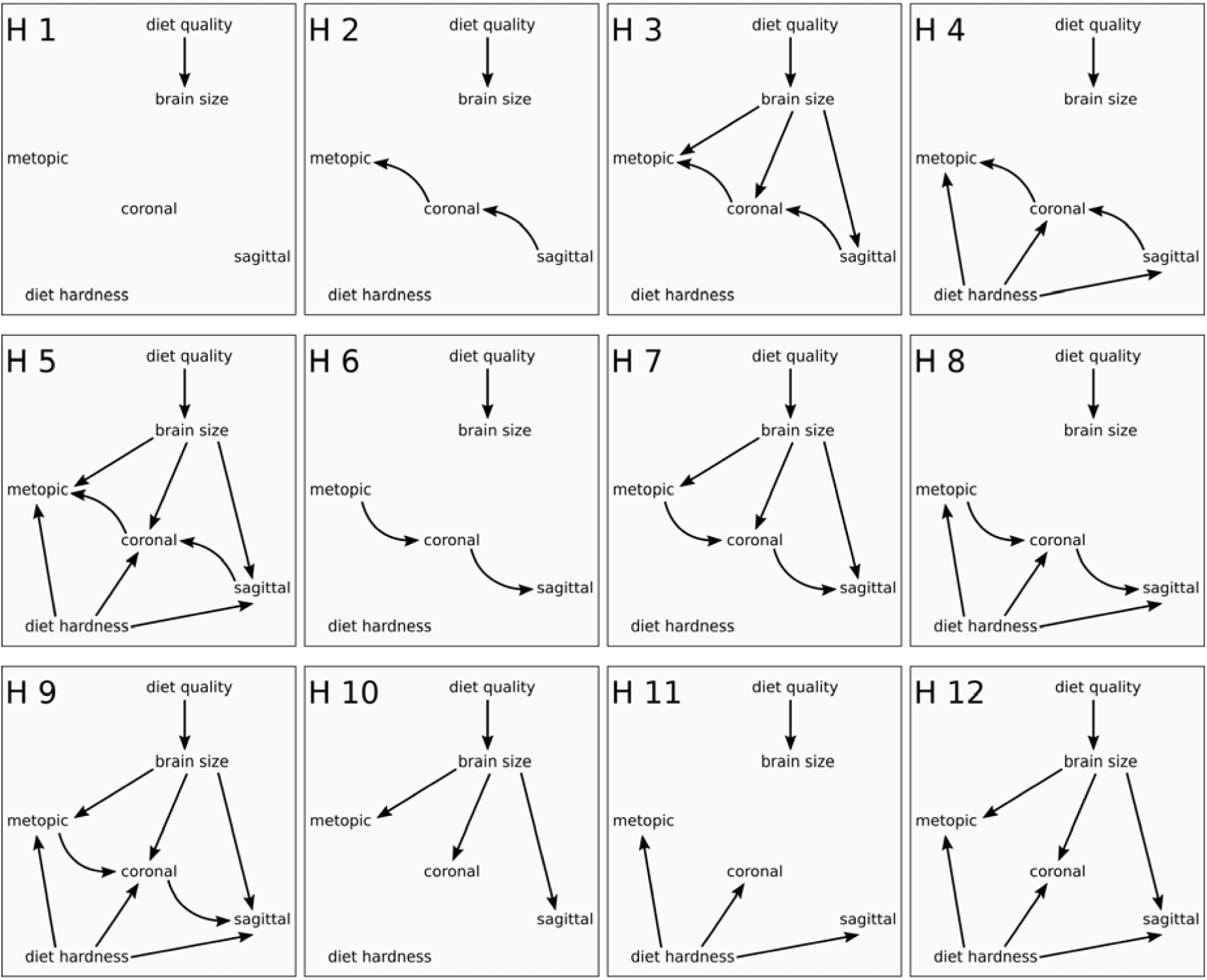
Alternative causal models compared with the phylogenetic path analysis. Model H1 is the null hypothesis of no causal relation among brain size, diet hardness, and sutures patency. H2 to H5 evaluate the causation of brain and/or diet when suture closure has a causal directionality following the relative order of closure in mammals (Rager et al., 2014). H6 to H9 evaluate the causation of brain and/or diet when suture closure follows the anteroposterior timing of ossification of the skull (Koyabu et al., 2014). H10 to H12 evaluate the influence of brain and diet in the absence of any type of developmental causation between sutures.

Brain and body mass information was gathered from a recent study on brain size allometry in mammals (Burger et al., 2019). We noticed an error in the body mass for *Bos taurus* (46,100 g) and fixed using the correct value (461,000 g) from the original reference (Isler & van Schaik, 2012). We favor Burger’s dataset because measurements were systematically compiled (e.g., two cross-reference check to assess authenticity of measurements, female-male averages except for dimorphic species), and brain and body mass for each species come from the same study, which minimizes potential errors. Brain size was calculated as the residual of a phylogenetic generalized least square regression of the log-transformation of brain mass against body mass.

Diet information was extracted from EltonTraits 1.0 database on species-level foraging attributes (Wilman et al., 2014). Data includes the percentage of the type of food consumed for each species. Diet quality was measured using Sailer and colleagues’ equation (Sailer et al., 1985) as, *DQ* = *plants* + 2 X (*fruit* + *seed* + *nectar*) + 3.5 X (*meat*), ranging from 100 to 350. Because there is not a similar measure for diet hardness based on the relative amount of food consumed, we followed an approach similar to that used for diet quality, measuring diet hardness as, *DH* = (*fruit* + *meat*) +2 X (*plants* + *invertebrates*) + 3.5 X (*seed* + *scanveing*), ranging from 0 to 350 (however, only species with a nectar-based diet will rank between 0 and 100). This relative division of food types by hardness (i.e., 1x, 2x, 3.5x) agrees with the division of hard foods used in experimental studies (Marcé-Nogué et al., 2017; Williams et al., 2005). Diet quality and hardness were both normalized between 0 and 1.

### Convergent amino acid substitutions (CAAS)

Multiple sequence alignments (MSA) were downloaded from the University of California Santa Cruz (UCSC) Genome Browser (Kent et al., 2002). We kept the 18533 MSA corresponding to the longest transcript of each gene. We then filtered out those sequences having more than 30% gaps or ambiguous amino acid definitions in any of the species analyzed. The final background pool of genes included 10922 MSA. On this gene pool, we searched for CAAS that co-occur in three pairs of closely related species with an opposite suture patency (open/closed) for the metopic, coronal, and sagittal sutures (**Figure 5**), for a total of nine pair-wise comparisons. Using an in-house script from a past study (Muntané et al., 2018), we retrieved all positions in which an amino acid differed between species with the suture open and species with the suture closed. We kept those cases in which an amino acid differed between the two groups and was shared by all the species of at least one group, discarding any case with a gap in that position. The final list of candidate genes includes only those genes that had convergent amino acid substitutions in the compared pairs for the three sutures.

**Figure 5.**
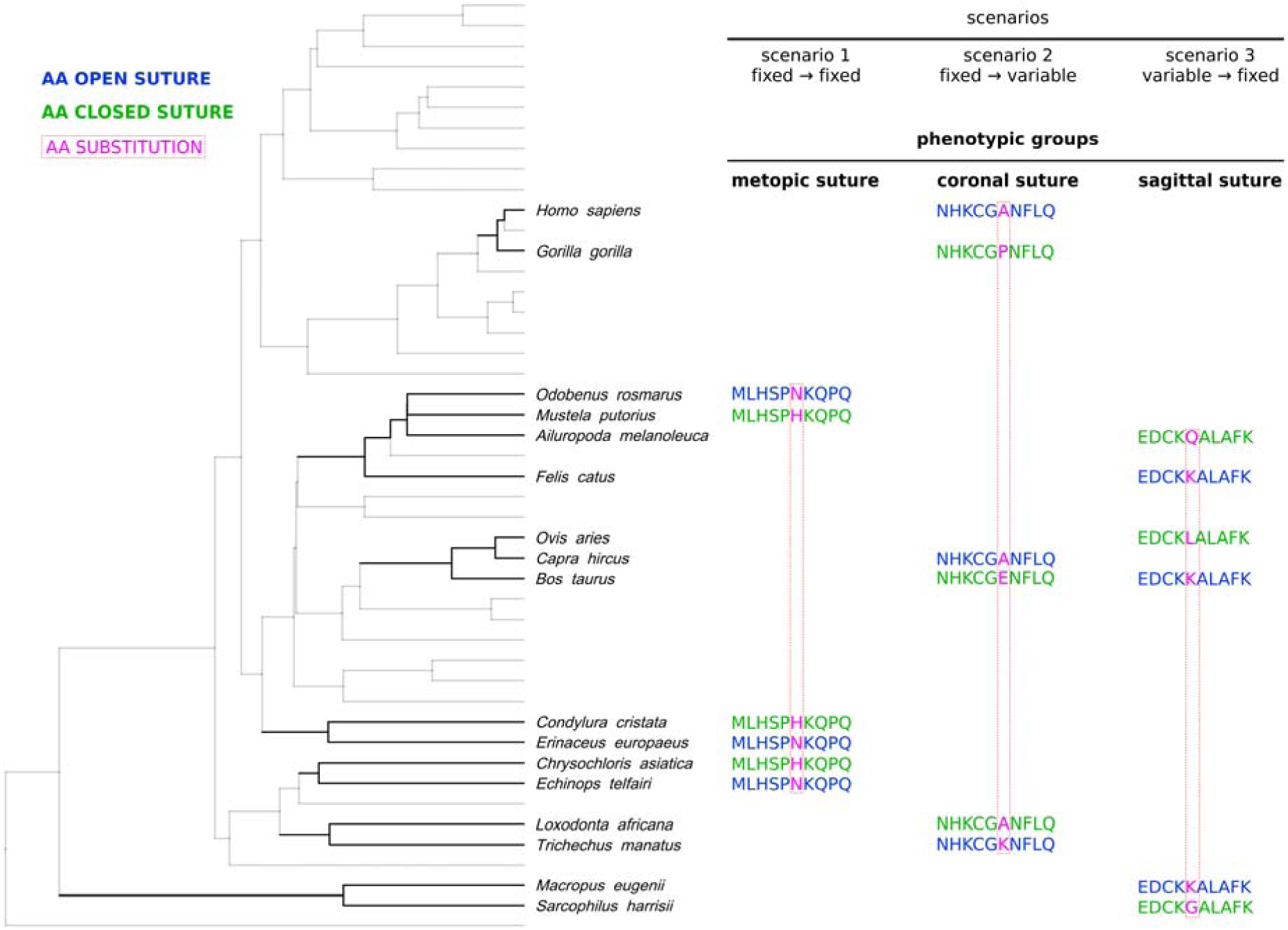
Structure of the convergent amino acid substitution analysis. Pairs of closely related species were selected for comparison based on their suture status as either open (blue) or closed (green), producing three phenotypic groups with three pairs of species each. Three scenarios of substitution were searched for every phenotypic group (only one is shown here for each suture).

We considered three scenarios or types of convergent substitution. Scenario 1 captures the same, single amino acid substitution for all pairs compared between species with the suture open and closed (e.g., open = {asparagine} → closed = {histidine}). Scenario 2 captures substitutions of a same fixed amino acid in species with the suture open to a variable set of different amino acids in species with the suture closed (e.g., open = {alanine} → closed = {proline, glutamate, lysine}). Scenario 3 is the reverse case of scenario 2: a variable set of amino acids in species with the suture open changed to a same amino acid in species with the suture closed (e.g., open = {glutamine, leucine, glycine} → closed = {lysine}).

### Statistical and internal validation of candidate genes

We performed a statistical validation of candidate genes using bootstrapping to assess whether the number of genes carrying CAAS were different than expected at random. For 1000 iterations, we sorted the 17 species analyzed (see **Figure 6**) into two random groups and scanned the background pool of genes for genes carrying at least one non-gapped CAAS. The bootstrap results span from 3,274 to 10,681 hits with a median of 8,898 hits, and 5% and 95% intervals are 5,759.8 and 10,082 genes hit, respectively. Finally, we tested the significance of the genes overlapping for the three sutures in R using the *SuperExactTest* package (Wang et al., 2015).

**Figure 6.**
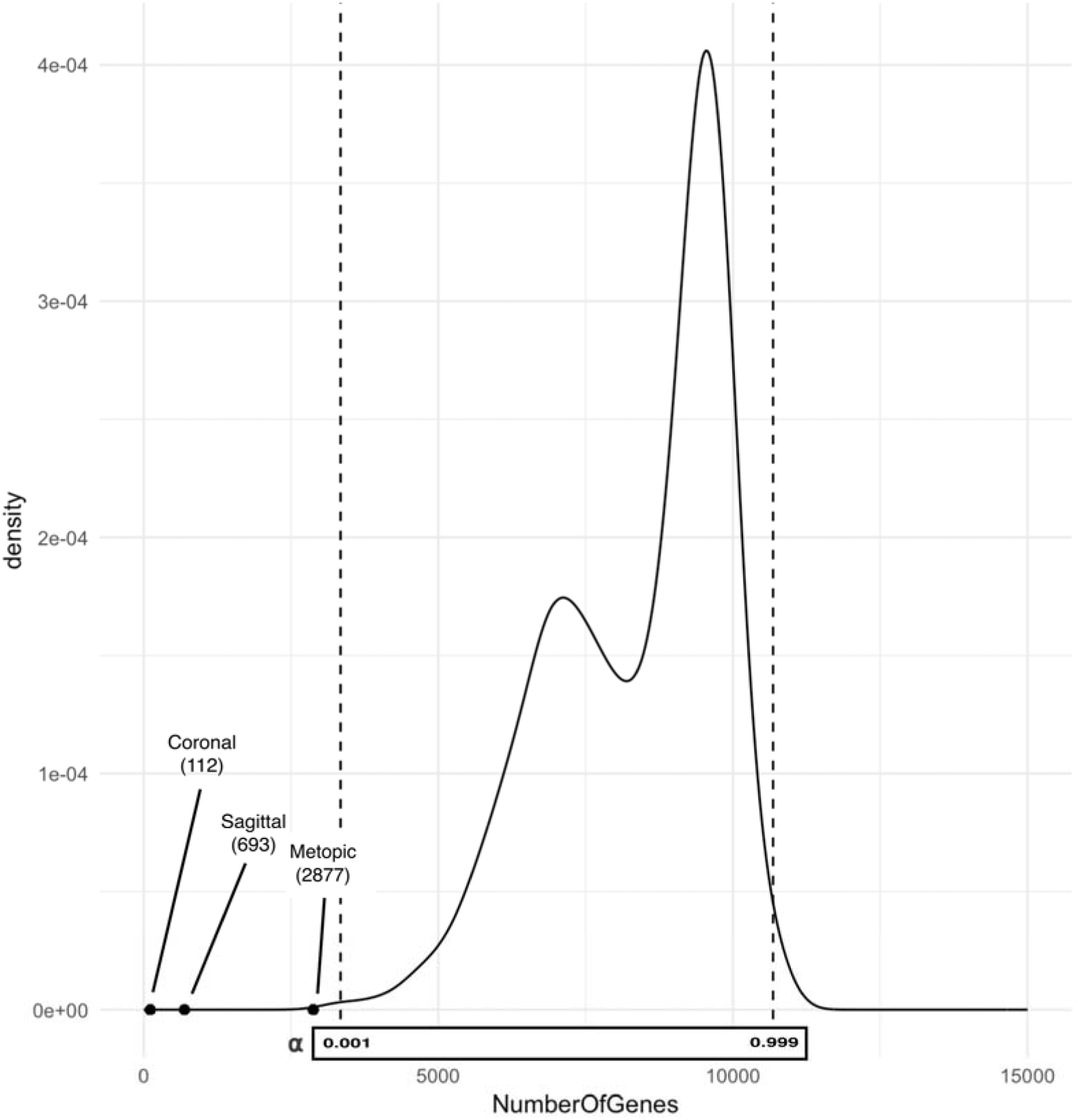
Results of the bootstrap comparisons of CAAS. Red lines show the 0.001 and 0.999 probability limits. Dots mark the total number of genes identified for each suture.

To assess biological significance, we performed an internal validation of candidate genes within the whole sample of mammals, following the same procedure described before. We considered a candidate gene as validated when it is also identified carrying open-versus-closed CAAS in the whole sample. Validated genes are discussed in the context of suture biology in more detail.

### Functional and pathological enrichment

We performed set enrichment analyses for the list of candidate gens using GOATOOLS (Klopfenstein et al., 2018) with the background pool of 10,922 protein-coding genes as reference. First, we performed a functional enrichment for biological processes, cellular components, and molecular functions of the Gene Ontology (GO) database (Ashburner et al., 2000; The Gene Ontology Consortium, 2019) using in-house scripts. Enrichments are based on GO definitions present in the go-basic.obo file available on Gene Ontology public database (Accessed February 2020). Then, we performed an enrichment analysis of candidate genes for three sets of genes associated to premature cranial suture closure (craniosynostosis) using Fisher’s exact test. The first set comprises 97 genes linked to (mostly) syndromic craniosynostosis conditions in the Human Phenotype Ontology (HP:0001363). The second set comprises 959 genes with differential gene expression profiles in RNA-Seq data for human osteoblast cultures derived from bone biopsy of nonsyndromic craniosynostosis cases (Rojas-Peña et al., 2014). The third set comprises 53 genes from two GWAS studies on nonsyndromic craniosynostosis: one for the sagittal suture (Justice et al., 2012) and one for the metopic suture (Justice et al., 2020). Note that we only included genes that are also present in our background pool of genes.

### Methodological limitations

#### Suture patency and sample size

The number of specimens examined of each species is uneven, and for some species only a few individuals were available. This is either because the species is rare and was not available in the natural history museums visited or in the online repositories consulted or because the covid-19 outbreak prevented us visiting additional museum collections. However, the conserved nature of suture patency gave us confidence that the suture patency measurements represent the general, highly conserved pattern of each species, and that its categorization as open or closed is valid. A broad sampling would provide stronger support to our conclusions. Moreover, it would make possible to include intraspecific variation within the analysis, which was not possible now because some species (perhaps due to the small sample size) showed little variation.

#### Quality of referenced genomes, alignments, and number of gaps

The quality of the mammalian reference genomes available for comparison is also uneven; specially compared to human and some model organisms. The consequence is that some alignments of protein-coding genes have a high number of gaps (i.e., not comparable positions because of unknown amino acid, deletion/insertion). These gapped positions complicate comparing simultaneously many species. In this study, we decided to take the most conservative approach: we excluded genes with more than 30% of positions with a gap from the background pool and also positions with gaps for each pairwise CAAS. The side effect of this strategy is that some genes are inevitably excluded because they show many gaps, among them genes linked to craniofacial development, such as ALX4, ERF, SMAD6. In addition, protein lengths may exert some bias in the genes excluded as well as the number of CAAS. Nevertheless, we adopted this conservative approach because the laxer alternative added an additional layer of ambiguity in the results. We hope that soon more and better referenced genomes will be available that allow expanding our comparative study.

## ACKNOWLDEGEMENTS

This work will not have been possible without the help from many colleagues that shared their museum skull photographs and CT scans, which mitigated the impossibility to visit museum collections due to COVID-19 lockdowns worldwide. We thank Borja Figueirido, Georgina Hutton, Jordi Marcé-Nogué, Laura O. Wilson, Mathew J. Stewart, Patrick Arnold, and Soledad De Esteban Trivigno. Also, to Christopher Smith for his photographs from the American Museum of Natural History, New York. We also thank curators Javier Quesada (Museum of Zoology of Barcelona), Ángel Luís Garvía Rodriguez and Luís Castelo Vicente (National Museum of Natural Sciences) for their help in accessing specimens. Finally, we are thankful to Tomàs Marquès-Bonet and his group, specially David Juan, for their insights and feedback.

## FUNDING

BE-A has received financial support for this project by the Postdoctoral Junior Leader Fellowship Programme from “la Caixa” Banking Foundation (LCF/BQ/LI18/11630002). BE-A and AN are supported by AEI-PGC2018-101927-BI00 (FEDER/UE). We also thank the Unidad de Excelencia María de Maeztu funded by the AEI (CEX2018-000792-M).

## COMPETING INTERESTS

Authors declare no competing interests.

## DATA AND CODE AVAILABILITY

Data and code used in this manuscript is available at https://figshare.com/projects/Cranial_Suture_Closure/81209

## AUTHOR CONTRIBUTIONS

BE-A designed the study, collected the data, and wrote the manuscript.

BE-A and FB analyzed the data.

FB, XF, GM, and AN conceived and designed the in-house scripts for CAAS. All authors interpreted the results and wrote the manuscript.

## Notes

https://figshare.com/projects/Cranial_Suture_Closure/81209

### Competing Interest Statement

The authors have declared no competing interest.

## REFERENCES

Adhikari, K., Fuentes-Guajardo, M., Quinto-Sánchez, M., Mendoza-Revilla, J., Camilo Chacón-Duque, J., Acuña-Alonzo, V., Jaramillo, C., Arias, W., Lozano, R. B., Pérez, G. M., Gómez-Valdés, J., Villamil-Ramírez, H., Hunemeier, T., Ramallo, V., Silva de Cerqueira, C. C., Hurtado, M., Villegas, V., Granja, V., Gallo, C., … Ruiz-Linares, A. (2016). A genome-wide association scan implicates *DCHS2, RUNX2, GLI3, PAX1* and *EDAR* in human facial variation. Nature Communications, 7, 11616. https://doi.org/10.1038/ncomms11616

Aiello, L. C., & Wheeler, P. (1995). The expensive-tissue hypothesis: The brain and the digestive system in human and primate evolution. Current Anthropology, 36(2), 199–221. https://doi.org/10.1086/204350.

Alföldi, J., & Lindblad-Toh, K. (2013). Comparative genomics as a tool to understand evolution and disease. Genome Research, 23(7), 1063–1068. https://doi.org/10.1101/gr.157503.113

Allen, K. L., & Kay, R. F. (2012). Dietary quality and encephalization in platyrrhine primates. Proceedings of the Royal Society B: Biological Sciences, 279(1729), 715–721. https://doi.org/10.1098/rspb.2011.1311

Al-Rekabi, Z., Cunningham, M. L., & Sniadecki, N. J. (2017). Cell mechanics of craniosynostosis. ACS Biomaterials Science & Engineering, 3(11), 2733–2743. https://doi.org/10.1021/acsbiomaterials.6b00557

Armand, T., Schaefer, E., Di Rocco, F., Edery, P., Collet, C., & Rossi, M. (2019). Genetic bases of craniosynostoses: An update. Neuro-Chirurgie, 65(5), 196–201. https://doi.org/10.1016/j.neuchi.2019.10.003

Ashburner, M., Ball, C. A., Blake, J. A., Botstein, D., Butler, H., Cherry, J. M., & Davis, A. P. (2000). Gene Ontology: Tool for the unification of biology. Nature Genetics, 25(1), 25–29. https://doi.org/10.1038/75556.

Atsawasuwan, P., Lu, X., Ito, Y., Zhang, Y., Evans, C. A., & Luan, X. (2013). Ameloblastin inhibits cranial suture closure by modulating *Msx2* expression and proliferation. PLOS ONE, 8(4), e52800. https://doi.org/10.1371/journal.pone.0052800

Barbeito-Andrés, J., Bonfili, N., Nogué, J. M., Bernal, V., & Gonzalez, P. N. (2020). Modeling the effect of brain growth on cranial bones using finite-element analysis and geometric morphometrics. Surgical and Radiologic Anatomy. https://doi.org/10.1007/s00276-020-02466-y

Bärmann, E. V., & Sánchez-Villagra, M. R. (2012). A phylogenetic study of late growth events in a mammalian evolutionary radiation: The cranial sutures of terrestrial artiodactyl mammals. Journal of Mammalian Evolution, 19(1), 43–56. https://doi.org/10.1007/s10914-011-9176-8

Bijl, W. van der. (2018). Phylopath: Easy phylogenetic path analysis in R. PeerJ, 6, e4718. https://doi.org/10.7717/peerj.4718

Bright, J. A. (2012). The importance of craniofacial sutures in biomechanical finite element models of the domestic pig. PLOS ONE, 7(2), e31769. https://doi.org/10.1371/journal.pone.0031769

Burger, J. R., George, M. A., Leadbetter, C., & Shaikh, F. (2019). The allometry of brain size in mammals. Journal of Mammalogy, 100(2), 276–283. https://doi.org/10.1093/jmammal/gyz043

Byron, C. D., Borke, J., Yu, J., Pashley, D., Wingard, C. J., & Hamrick, M. (2004). Effects of increased muscle mass on mouse sagittal suture morphology and mechanics. The Anatomical Record Part A: Discoveries in Molecular, Cellular, and Evolutionary Biology, 279A(1), 676–684. https://doi.org/10.1002/ar.a.20055

Cohen, M., & MacLean, R. (2000). Craniosynostosis: Diagnosis, evaluation, and management. Oxford University Press.

Cornille, M., Dambroise, E., Komla-Ebri, D., Kaci, N., Biosse-Duplan, M., Rocco, F. D., & Legeai-Mallet, L. (2019). Animal models of craniosynostosis. Neurochirurgie, 65(5), 202–209. https://doi.org/10.1016/j.neuchi.2019.09.010

Cray, J., Cooper, G. M., Mooney, M. P., & Siegel, M. I. (2014). Ectocranial suture fusion in primates: Pattern and phylogeny. Journal of Morphology, 275(3), 342–347. https://doi.org/10.1002/jmor.20218

Cuellar, A., Bala, K., Di Pietro, L., Barba, M., Yagnik, G., Liu, J. L., Stevens, C., Hur, D. J., Ingersoll, R. G., Justice, C. M., Drissi, H., Kim, J., Lattanzi, W., & Boyadjiev, S. A. (2020). Gain-of-function variants and overexpression of RUNX2 in patients with nonsyndromic midline craniosynostosis. Bone, 115395. https://doi.org/10.1016/j.bone.2020.115395

Davis, A. A., Zuccoli, G., Haredy, M. M., Losee, J., Pollack, I. F., Madan-Khetarpal, S., Goldstein, J. A., & Nischal, K. K. (2019). RASopathy in patients with isolated sagittal synostosis. Global Pediatric Health. https://doi.org/10.1177/2333794x19846774

de Magalhães, J. P., & Wang, J. (2019). The fog of genetics: What is known, unknown and unknowable in the genetics of complex traits and diseases. EMBO Reports, 20(11), e48054. https://doi.org/10.15252/embr.201948054

Dempsey, R. F., Monson, L. A., Maricevich, R. S., Truong, T. A., Olarunnipa, S., Lam, S. K., Dauser, R. C., Hollier, L. H., & Buchanan, E. P. (2019). Nonsyndromic craniosynostosis. Clinics in Plastic Surgery, 46(2), 123–139. https://doi.org/10.1016/j.cps.2018.11.001

Esteve-Altava, B., & Rasskin-Gutman, D. (2014). Beyond the functional matrix hypothesis: A network null model of human skull growth for the formation of bone articulations. Journal of Anatomy, 225(3), 306–316. https://doi.org/10.1111/joa.12212

Esteve-Altava, B., & Rasskin-Gutman, D. (2015). Evo-Devo insights from pathological networks: Exploring craniosynostosis as a developmental mechanism for modularity and complexity in the human skull. Journal of Anthropological Sciences, 93, 103–117. https://doi.org/10.4436/JASS.93001

Esteve-Altava, B., Vallès-Català, T., Guimerà, R., Sales-Pardo, M., & Rasskin-Gutman, D. (2017). Bone fusion in normal and pathological development is constrained by the network architecture of the human skull. Scientific Reports, 7(1), 3376. https://doi.org/10.1038/s41598-017-03196-9

Falk, D., Zollikofer, C. P. E., Morimoto, N., & Ponce de Leon, M. S. (2012). Metopic suture of Taung (*Australopithecus africanus*) and its implications for hominin brain evolution. Proceedings of the National Academy of Sciences, 109(22), 8467–8470. https://doi.org/10.1073/pnas.1119752109

Foote, A. D., Liu, Y., Thomas, G. W. C., Vinař, T., Alföldi, J., Deng, J., Dugan, S., van Elk, C. E., Hunter, M. E., Joshi, V., Khan, Z., Kovar, C., Lee, S. L., Lindblad-Toh, K., Mancia, A., Nielsen, R., Qin, X., Qu, J., Raney, B. J., … Gibbs, R. A. (2015). Convergent evolution of the genomes of marine mammals. Nature Genetics, 47(3), 272–275. https://doi.org/10.1038/ng.3198

Garza, R. M., & Khosla, R. K. (2012). Nonsyndromic craniosynostosis. Seminars in Plastic Surgery, 26(2), 53–63. https://doi.org/10.1055/s-0032-1320063

Green, R. E., Krause, J., Briggs, A. W., Maricic, T., Stenzel, U., Kircher, M., Patterson, N., Li, H., Zhai, W., Fritz, M. H.-Y., Hansen, N. F., Durand, E. Y., Malaspinas, A.-S., Jensen, J. D., Marques-Bonet, T., Alkan, C., Prüfer, K., Meyer, M., Burbano, H. A., … Pääbo, S. (2010). A draft sequence of the Neandertal genome. Science (New York, N.Y.), 328(5979), 710–722. https://doi.org/10.1126/science.1188021

Grova, M., Lo, D. D., Montoro, D., Hyun, J. S., Chung, M. T., Wan, D. C., & Longaker, M. T. (2012). Animal models of cranial suture biology. The Journal of Craniofacial Surgery, 23(7 0 1), 1954–1958. https://doi.org/10.1097/SCS.0b013e318258ba53

Herring, S. W. (2008). Mechanical influences on suture development and patency. In D. P. Rice (Ed.), Craniofacial Sutures. Development, Disease and Treatment (Vol. 12, pp. 41–56). S. KARGER AG. https://doi.org/10.1159/000115031

Herring, & Teng, S. (2000). Strain in the braincase and its sutures during function. American Journal of Physical Anthropology, 112(4), 575–593. https://doi.org/10.1002/1096-8644(200008)112:4<575::AID-AJPA10>3.0.CO;2-0

Imerslund, O. (1951). Craniostenosis and vitamin D resistant rickets. Acta Paediatrica, 40(5), 449–456. https://doi.org/10.1111/j.1651-2227.1951.tb16509.x

Isler, K., & van Schaik, C. P. (2012). Allomaternal care, life history and brain size evolution in mammals. Journal of Human Evolution, 63(1), 52–63. https://doi.org/10.1016/j.jhevol.2012.03.009

Jaszczuk, P., Rogers, G. F., Guzman, R., & Proctor, M. R. (2016). X-linked hypophosphatemic rickets and sagittal craniosynostosis: Three patients requiring operative cranial expansion: case series and literature review. Child’s Nervous System, 32(5), 887–891. https://doi.org/10.1007/s00381-015-2934-9

Justice, C. M., Cuellar, A., Bala, K., Sabourin, J. A., Cunningham, M. L., Crawford, K., Phipps, J. M., Zhou, Y., Cilliers, D., Byren, J. C., Johnson, D., Wall, S. A., Morton, J. E. V., Noons, P., Sweeney, E., Weber, A., Rees, K. E. M., Wilson, L. C., Simeonov, E., … National Birth Defects Prevention Study. (2020). A genome-wide association study implicates the *BMP7* locus as a risk factor for nonsyndromic metopic craniosynostosis. Human Genetics. https://doi.org/10.1007/s00439-020-02157-z

Justice, C. M., Yagnik, G., Kim, Y., Peter, I., Jabs, E. W., Erazo, M., Ye, X., Ainehsazan, E., Shi, L., Cunningham, M. L., Kimonis, V., Roscioli, T., Wall, S. A., Wilkie, A. O. M., Stoler, J., Richtsmeier, J. T., Heuzé, Y., Sanchez-Lara, P. A., Buckley, M. F., … Boyadjiev, S. A. (2012). A genome-wide association study identifies susceptibility loci for nonsyndromic sagittal craniosynostosis near *BMP2* and within *BBS9*. Nature Genetics, 44(12), 1360–1364. https://doi.org/10.1038/ng.2463

Katsianou, M. A., Adamopoulos, C., Vastardis, H., & Basdra, E. K. (2016). Signaling mechanisms implicated in cranial sutures pathophysiology: Craniosynostosis. BBA Clinical, 6, 165–176. https://doi.org/10.1016/j.bbacli.2016.04.006

Kent, W. J., Sugnet, C. W., Furey, T. S., Roskin, K. M., Pringle, T. H., Zahler, A. M., & Haussler, and D. (2002). The human genome browser at UCSC. Genome Research, 12(6), 996–1006. https://doi.org/10.1101/gr.229102

Khonsari, R. H., Olivier J., Vigneaux P., Sanchez S., Tafforeau P., Ahlberg P. E., Di Rocco F., Bresch D., Corre P., Ohazama A., Sharpe P. T., & Calvez V. (2013). A mathematical model for mechanotransduction at the early steps of suture formation. Proceedings of the Royal Society B, 280(1759), 20122670. https://doi.org/10.1098/rspb.2012.2670

Klopfenstein, D. V., Zhang, L., Pedersen, B. S., Ramírez, F., Warwick Vesztrocy, A., Naldi, A., Mungall, C. J., Yunes, J. M., Botvinnik, O., Weigel, M., Dampier, W., Dessimoz, C., Flick, P., & Tang, H. (2018). GOATOOLS: A Python library for Gene Ontology analyses. Scientific Reports, 8(1), 10872. https://doi.org/10.1038/s41598-018-28948-z

Koyabu, D., Endo, H., Mitgutsch, C., Suwa, G., Catania, K. C., Zollikofer, C. P., Oda, S., Koyasu, K., Ando, M., & Sánchez-Villagra, M. R. (2011). Heterochrony and developmental modularity of cranial osteogenesis in lipotyphlan mammals. EvoDevo, 2(1), 21. https://doi.org/10.1186/2041-9139-2-21

Koyabu, D., Werneburg, I., Morimoto, N., Zollikofer, C. P. E., Forasiepi, A. M., Endo, H., Kimura, J., Ohdachi, S. D., Truong Son, N., & Sánchez-Villagra, M. R. (2014). Mammalian skull heterochrony reveals modular evolution and a link between cranial development and brain size. Nature Communications, 5(1). https://doi.org/10.1038/ncomms4625

Lattanzi, W., Barba, M., Di Pietro, L., & Boyadjiev, S. A. (2017). Genetic advances in craniosynostosis. American Journal of Medical Genetics, 173(5), 1406–1429. https://doi.org/10.1002/ajmg.a.38159

Lieberman, D. E. (2010). The Evolution of the Human Head. Belknap Press Harvard University Press.

Lieberman, D. E. (2011). Epigenetic integration, complexity, and evolvability of the head. In B. Hallgrímsson & B. K. Hall (Eds.), Epigenetics. Linking Genotype and Phenotype in Development and Evolution (pp. 271–289). University of California Press.

Maeno, T., Moriishi, T., Yoshida, C. A., Komori, H., Kanatani, N., Izumi, S., Takaoka, K., & Komori, T. (2011). Early onset of *Runx2* expression caused craniosynostosis, ectopic bone formation, and limb defects. Bone, 49(4), 673–682. https://doi.org/10.1016/j.bone.2011.07.023

Magherini, S., Fiore, M. G., Chiarelli, B., Serrao, A., Paternostro, F., Morucci, G., Branca, J. J. V., Ruggiero, M., & Pacini, S. (2015). Metopic suture and RUNX2, a key transcription factor in osseous morphogenesis with possible important implications for human brain evolution. Italian Journal of Anatomy and Embryology, 120(1), 5–20.

Marcé-Nogué, J., Püschel, T. A., & Kaiser, T. M. (2017). A biomechanical approach to understand the ecomorphological relationship between primate mandibles and diet. Scientific Reports, 7(1), 1–12. https://doi.org/10.1038/s41598-017-08161-0

Moazen, M., Curtis, N., O’Higgins, P., Jones, M. E. H., Evans, S. E., & Fagan, M. J. (2009). Assessment of the role of sutures in a lizard skull: A computer modelling study. Proceedings of the Royal Society B: Biological Sciences, 276(1654), 39–46. https://doi.org/10.1098/rspb.2008.0863

Morriss-Kay, G. M., & Wilkie, A. O. M. (2005). Growth of the normal skull vault and its alteration in craniosynostosis: Insights from human genetics and experimental studies. Journal of Anatomy, 207(5), 637–653. https://doi.org/10.1111/j.1469-7580.2005.00475.x

Moss, M. L. (1975). Functional anatomy of cranial synostosis. Child’s Brain, 1(1), 22–33.

Moss, M. L., & Young, R. W. (1960). A functional approach to craniology. American Journal of Physical Anthropology, 18(4), 281–292.

Muntané, G., Farré, X., Rodríguez, J. A., Pegueroles, C., Hughes, D. A., de Magalhães, J. P., Gabaldón, T., & Navarro, A. (2018). Biological processes modulating longevity across primates: A phylogenetic genome-phenome analysis. Molecular Biology and Evolution, 35(8), 1990–2004. https://doi.org/10.1093/molbev/msy105

Newton, A. H., Feigin, C. Y., & Pask, A. J. (2017). *RUNX2* repeat variation does not drive craniofacial diversity in marsupials. BMC Evolutionary Biology, 17(1). https://doi.org/10.1186/s12862-017-0955-6

Oh, J., Kim, Y. K., Yasuda, M., Koyabu, D., & Kimura, J. (2017). Cranial suture closure pattern in water deer and implications of suture evolution in cervids. Mammalian Biology, 86, 17–20. https://doi.org/10.1016/j.mambio.2017.03.004

Opperman, L. A. (2000). Cranial sutures as intramembranous bone growth sites. Developmental Dynamics, 219, 472–485.

Opperman, L. A., & Rawlins, J. T. (2005). The extracellular matrix environment in suture morphogenesis and growth. Cells, Tissues, Organs, 181 (3–4), 127–135. https://doi.org/10.1159/000091374

Payne, J. L., & Wagner, A. (2019). The causes of evolvability and their evolution. Nature Reviews Genetics, 20(1), 24–38. https://doi.org/10.1038/s41576-018-0069-z

Poot, M. (2019). Structural genome variations related to craniosynostosis. Molecular Syndromology, 10(1–2), 24–39. https://doi.org/10.1159/000490480

R Core Team. (2019). R: A language and environment for statistical computing (Version 3.6.2) [Computer software]. R Foundation for Statistical Computing. https://www.R-project.org/

Rafferty, K. L., Baldwin, M. C., Soh, S. H., & Herring, S. W. (2019). Mechanobiology of bone and suture—Results from a pig model. Orthodontics & Craniofacial Research, 22 Suppl 1, 82–89. https://doi.org/10.1111/ocr.12276

Rafferty, K. L., & Herring, S. W. (1999). Craniofacial sutures: Morphology, growth, and in vivo masticatory strains. Journal of Morphology, 242(2), 167–179. https://doi.org/10.1002/(SICI)1097-4687(199911)242:2<167::AID-JMOR8>3.0.CO;2-1

Rager, L., Hautier, L., Forasiepi, A., Goswami, A., & Sánchez-Villagra, M. R. (2014). Timing of cranial suture closure in placental mammals: Phylogenetic patterns, intraspecific variation, and comparison with marsupials. Journal of Morphology, 275(2), 125–140. https://doi.org/10.1002/jmor.20203

Rasskin-Gutman, D., & Esteve-Altava, B. (2018). Concept of burden in evo-devo. In L. Nuno de la Rosa & G. Müller (Eds.), Evolutionary Developmental Biology: A Reference Guide (pp. 1–11). Springer International Publishing.

Richtsmeier, J. T. (2018). A century of development. American Journal of Physical Anthropology, 165(4), 726–740. https://doi.org/10.1002/ajpa.23379

Richtsmeier, J. T., Aldridge, K., DeLeon, V. B., Panchal, J., Kane, A. A., Marsh, J. L., Yan, P., & Cole, T. M. (2006). Phenotypic integration of neurocranium and brain. Journal of Experimental Zoology Part B: Molecular and Developmental Evolution, 306B (4), 360–378. https://doi.org/10.1002/jez.b.21092

Richtsmeier, J. T., & Flaherty, K. (2013). Hand in glove: Brain and skull in development and dysmorphogenesis. Acta Neuropathologica, 125(4), 469–489. https://doi.org/10.1007/s00401-013-1104-y

Rojas-Peña, M. L., Olivares-Navarrete, R., Hyzy, S., Arafat, D., Schwartz, Z., Boyan, B. D., Williams, J., & Gibson, G. (2014). Characterization of distinct classes of differential gene expression in osteoblast cultures from non-syndromic craniosynostosis bone. Journal of Genomics, 2, 121–130. https://doi.org/10.7150/jgen.8833

Roosenboom, J., Lee, M. K., Hecht, J. T., Heike, C. L., Wehby, G. L., Christensen, K., Feingold, E., Marazita, M. L., Maga, A. M., Shaffer, J. R., & Weinberg, S. M. (2018). Mapping genetic variants for cranial vault shape in humans. PLOS ONE, 13(4), e0196148. https://doi.org/10.1371/journal.pone.0196148

Roston, R. A., & Roth, V. L. (2019). Cetacean skull telescoping brings evolution of cranial sutures into focus. The Anatomical Record, 302(7), 1055–1073. https://doi.org/10.1002/ar.24079

Rymer, K. (2015). Identification of Candidate Genes for Craniosynostosis [Virginia Commonwealth University]. https://scholarscompass.vcu.edu/etd/3782

Sailer, L. D., Gaulin, S. J. C., Boster, J. S., & Kurland, J. A. (1985). Measuring the relationship between dietary quality and body size in primates. Primates, 26(1), 14–27. https://doi.org/10.1007/BF02389044

Salpea, P., Russanova, V. R., Hirai, T. H., Sourlingas, T. G., Sekeri-Pataryas, K. E., Romero, R., Epstein, J., & Howard, B. H. (2012). Postnatal development- and age-related changes in DNA-methylation patterns in the human genome. Nucleic Acids Research, 40(14), 6477–6494. https://doi.org/10.1093/nar/gks312

Sewda, A., White, S. R., Erazo, M., Hao, K., García-Fructuoso, G., Fernández-Rodriguez, I., Heuzé, Y., Richtsmeier, J. T., Romitti, P. A., Reva, B., Jabs, E. W., & Peter, I. (2019). Nonsyndromic craniosynostosis: Novel coding variants. Pediatric Research, 85(4), 463–468. https://doi.org/10.1038/s41390-019-0274-2

Smith, S. D., Pennell, M. W., Dunn, C. W., & Edwards, S. V. (2020). Phylogenetics is the new genetics (for most of biodiversity). Trends in Ecology & Evolution, 35(5), 415–425. https://doi.org/10.1016/j.tree.2020.01.005

Spector, J. A., Greenwald, J. A., Warren, S. M., Bouletreau, P. J., Crisera, F. E., Mehrara, B. J., & Longaker, M. T. (2002). Co-culture of osteoblasts with immature dural cells causes an increased rate and degree of osteoblast differentiation. Plastic and Reconstructive Surgery, 109(2), 631–642. https://doi.org/10.1097/00006534-200202000-00033

Stamper, B. D., Park, S. S., Beyer, R. P., Bammler, T. K., Farin, F. M., Mecham, B., & Cunningham, M. L. (2011). Differential expression of extracellular matrix-mediated pathways in single-suture craniosynostosis. PLOS ONE, 6(10). https://doi.org/10.1371/journal.pone.0026557

Sun, Z., Lee, E., & Herring, S. W. (2004). Cranial sutures and bones: Growth and fusion in relation to masticatory strain. The Anatomical Record, 276(2), 150–161. https://doi.org/10.1002/ar.a.20002

The Gene Ontology Consortium. (2019). The Gene Ontology Resource: 20 years and still GOing strong. Nucleic Acids Research, 47(D1), D330–D338. https://doi.org/10.1093/nar/gky1055

Tintut, Y., & Demer, L. L. (2014). Effects of bioactive lipids and lipoproteins on bone. Trends in Endocrinology and Metabolism, 25(2), 53–59. https://doi.org/10.1016/j.tem.2013.10.001

Twigg, S. R. F., Forecki, J., Goos, J. A. C., Richardson, I. C. A., Hoogeboom, A. J. M., van den Ouweland, A. M. W., Swagemakers, S. M. A., Lequin, M. H., Van Antwerp, D., McGowan, S. J., Westbury, I., Miller, K. A., Wall, S. A., WGS500 Consortium, van der Spek, P. J., Mathijssen, I. M. J., Pauws, E., Merzdorf, C. S., & Wilkie, A. O. M. (2015). Gain-of-function mutations in ZIC1 are associated with coronal craniosynostosis and learning disability. American Journal of Human Genetics, 97(3), 378–388. https://doi.org/10.1016/j.ajhg.2015.07.007

Twigg, S. R. F., & Wilkie, A. O. M. (2015). A genetic-pathophysiological framework for craniosynostosis. American Journal of Human Genetics, 97(3), 359–377. https://doi.org/10.1016/j.ajhg.2015.07.006

von Hardenberg, A., & Gonzalez[Voyer, A. (2013). Disentangling evolutionary cause-effect relationships with phylogenetic confirmatory path analysis. Evolution, 67(2), 378–387. https://doi.org/10.1111/j.1558-5646.2012.01790.x

Wang, M., Zhao, Y., & Zhang, B. (2015). Efficient test and visualization of multi-set intersections. Scientific Reports, 5(1), 16923. https://doi.org/10.1038/srep16923

Wang, P. I., Marcus, J. R., Fuchs, H. E., & Mukundan, S. (2015). Craniosynostosis secondary to rickets: Manifestations on computed tomography. Radiology Case Reports, 2(3). https://doi.org/10.2484/rcr.v2i3.43

Wilkie, A. O. M., Johnson, D., & Wall, S. A. (2017). Clinical genetics of craniosynostosis. Current Opinion in Pediatrics, 29(6), 622–628. https://doi.org/10.1097/MOP.0000000000000542

Williams, S. H., Wright, B. W., Truong, V. den, Daubert, C. R., & Vinyard, C. J. (2005). Mechanical properties of foods used in experimental studies of primate masticatory function. American Journal of Primatology, 67(3), 329–346. https://doi.org/10.1002/ajp.20189

Wilman, H., Belmaker, J., Simpson, J., Rosa, C. de la, Rivadeneira, M. M., & Jetz, W. (2014). EltonTraits 1.0: Species-level foraging attributes of the world’s birds and mammals. Ecology, 95(7), 2027–2027. https://doi.org/10.1890/13-1917.1

Wilson, L. A. B., & Sánchez Villagra, M. R. (2009). Heterochrony and patterns of cranial suture closure in hystricognath rodents. Journal of Anatomy, 214(3), 339–354. https://doi.org/10.1111/j.1469-7580.2008.01031.x

Wu, D.-D., Jin, W., Hao, X.-D., Tang, N. L. S., & Zhang, Y.-P. (2010). Evidence for positive selection on the osteogenin (BMP3) gene in human populations. PLOS ONE, 5(6), e10959. https://doi.org/10.1371/journal.pone.0010959

Wu, D.-D., Li, G.-M., Jin, W., Li, Y., & Zhang, Y.-P. (2012). Positive selection on the osteoarthritis-risk and decreased-height associated variants at the GDF5 Gene in East Asians. PLOS ONE, 7(8), e42553. https://doi.org/10.1371/journal.pone.0042553

Yamashita, S., Takeshima, H., Matsumoto, F., Yamasaki, K., Fukushima, T., Sakoda, H., Nakazato, M., Saito, K., Mizuguchi, A., Watanabe, T., Ohta, H., & Yokogami, K. (2019). Detection of the *KIAA1549*-*BRAF* fusion gene in cells forming microvascular proliferations in pilocytic astrocytoma. PLOS ONE, 14(7), e0220146. https://doi.org/10.1371/journal.pone.0220146

Zhou, X., Seim, I., & Gladyshev, V. N. (2015). Convergent evolution of marine mammals is associated with distinct substitutions in common genes. Scientific Reports, 5(1), 16550. https://doi.org/10.1038/srep16550

